# Antimycobacterial potential of Mycobacteriophage under disease-mimicking conditions

**DOI:** 10.1101/2020.05.16.100123

**Authors:** Yeswanth Chakravarthy Kalapala, Pallavi Raj Sharma, Rachit Agarwal

## Abstract

Antibiotic resistance continues to be a major global health risk with an increase in multi-drug resistant infections seen across nearly all bacterial diseases. Mycobacterial infections such as Tuberculosis (TB) and Non-Tuberculosis infections have seen a significant increase in the incidence of multi-drug resistant and extensively drug-resistant infections. With this increase in drug-resistant Mycobacteria, mycobacteriophage therapy offers a promising alternative. However, a comprehensive study on the infection dynamics of mycobacteriophage against their host bacteria and the evolution of bacteriophage (phage) resistance in the bacteria remains elusive. We aim to study the infection dynamics of a phage cocktail against Mycobacteria under various pathophysiological conditions such as low pH, low growth rate and hypoxia. We show that mycobacteriophages are effective against *M. smegmatis* under various conditions and the phage cocktail prevents emergence of resistance for long durations. Although the phages are able to amplify after infection, the initial multiplicity of infection plays an important role in reducing the bacterial growth and prolonging efficacy. Mycobacteriophages are effective against antibiotic-resistant strains of Mycobacterium and show synergy with antibiotics such as rifampicin and isoniazid. Finally, we also show that mycobacteriophages are efficient against *M. tuberculosis* both under lag and log phase for several weeks. These findings have important implications for developing phage therapy for Mycobacterium.

## 1 Introduction

Prolonged and unsupervised use of antibiotics results in an increase of antibiotic resistance across all bacterial species which is considered a major threat to human health (Keshavjee and Farmer, 2012; Sprenger and Fukuda, 2016). The Mycobacterium genus with over 190 recognized species contains various pathogenic species including *M. tuberculosis*, *M. leprae*, *M. abscessus*, *M. avium* complex, etc. (Rastogi et al., 2001). These infections are often difficult to treat owing to their resilient cell walls which render them resistant to prolonged exposure to acidic and alkaline environments (Rastogi et al., 2001). In addition, several mycobacterial species such as *M. leprae, M. abscessus* and *M. tuberculosis* rapidly evolve resistance to antibiotics (Iseman, 1994; Cambau et al., 2018; Luthra et al., 2018; Geneva: World Health Organisation, 2019). Similarly, the *Mycobacterium avium* Complex (MAC) infections are noted to be on the rise, especially in AIDS and pulmonary disease patients (Horsburgh, 1997) and MAC infections are often associated with an increase in mortality (Horsburgh, 1997; Diel et al., 2018). The antibiotic treatment regimen is associated with the distinct and long-lasting changes to the gut flora, often resulting in the depletion of various species of commensal bacteria (Wipperman et al., 2017). Antibiotics are also associated with various adverse side-effects ranging from nausea, vomiting and fever to anaphylaxis, hypersensitivity and photodermatitis (Castells et al., 2019). In order to tackle this growing rate of antibiotic resistance and the side effects of antibiotic treatment regimens, various therapeutic strategies have been proposed, including phage therapy, that has shown tremendous potential against various bacterial pathogens (Kortright et al., 2019).

Phages are viruses that are host specific towards bacteria and are inherently non-pathogenic to humans. They are abundantly present in the natural microflora of the human body (De Paepe et al., 2014; Manrique et al., 2016). A recent non-comparative clinical trial showed that the intravenous delivery of anti-staphylococcal phages in patients with severe infection did not pose any challenges to tolerability and safety (Petrovic Fabijan et al., 2020). Moreover, phages remain infective and lyse antibiotic-resistant bacteria (Kortright et al., 2019). To circumvent the narrow-host range specificity and development of resistance (Labrie et al., 2010), multiple phages are used in phage therapy. Successful compassionate use of phages against antibiotic-resistant bacteria on patients in the USA and Europe has been on the rise and shows their safety and translation potential (Jennes et al., 2017; Schooley et al., 2017).

With over 11,000 mycobacteriophages isolated and more than 1800 sequenced (Russell and Hatfull, 2017), there is a large library of phages against Mycobacterium. Although the desired lytic phenotype of phages (Pimentel, 2014; Catalão and Pimentel, 2018) for phage therapy brings the number down, development of Bacteriophage Recombineering of Electroporated DNA (BRED) technique (Marinelli et al., 2008) allows rapid conversion of lysogenic phages to lytic phages for phage therapy. Recently, a three-phage cocktail has been successfully used in a clinic for the treatment of a cystic fibrosis patient suffering from *Mycobacterium abscessus* infection (Dedrick et al., 2019). Prophylactic phage (D29) therapy against TB also showed some success in a preclinical animal model in reducing the burden of infection (Carrigy et al., 2019). A study employing phage therapy in treating subcutaneously injected *M. tuberculosis* (H37Rv) infection in guinea pigs showed that mycobacteriophages have a therapeutic effect (Sula et al., 1981).

Phage therapy is a strong candidate for treating mycobacterial infections, however it presents a few physiological hurdles that need to be taken into consideration. Various Mycobacterium species (*M. tuberculosis*, *M. avium* complex, *M. bovis* and *M. leprae*) are found to reside intracellularly at low pH conditions which poses a challenge in efficacy of anti-bacterials (Sibling et al., 1987; Oh and Straubinger, 1996; Ehlers and Schaible, 2012; Benítez-Guzmán et al., 2019). *M. tuberculosis* and *M. avium* complex also form granulomatous structures, marked by a gradient of hypoxia, nutrient deprivation and acidity (Timm et al., 2003; Voskuil et al., 2004; Lawn and Zumla, 2011; Ehlers and Schaible, 2012; Seto et al., 2020). For a phage to successfully eliminate these bacilli, it needs to be active and efficient under such low pH and hypoxic conditions and should be able to enter into the eukaryotic cells. Since most bacteria enter a non-replicative stationary phase, the phage should also be able to infect and kill bacteria in the stationary phase where most of the current antibiotics fail. Designing a phage therapy that would be effective in such complex physiological environments would necessitate a better understanding of the bacteria-phage dynamics under various conditions. However, to our knowledge, no study has looked at the efficacy of phages over time in such different environments. Hence, there is a need to study the efficacy of phage cocktails in the physiological and pathophysiological conditions in which bacteria is known to survive.

In this study, we have performed phage-bacteria kinetics under various pathophysiological conditions of mycobacterial diseases-low pH, hypoxia and stationary phase. We chose *M. smegmatis* as our model organism in designing the experiments owing to its relatively fast replication time and therefore, its ability to offer rapid screening and understanding of phage kinetics in extreme host environments. We study the emergence of phage resistance in *M. smegmatis* and the combinatory effect of antibiotics and phage therapy. We show that mycobacteriophage cocktails are effective against Mycobacterium under the low pH, hypoxic and stationary conditions and showed synergy with antibiotics. Finally, we also find that mycobacteriophages are able to prevent growth of *Mycobacterium tuberculosis* which encourages further exploration of phages for diseases caused by Mycobacterium.

## 2 Materials and Methods

### 2.1 Chemicals and Reagents

Middlebrook 7H9 Broth, Middlebrook 7H10 Agar and DMSO were purchased from Sigma (St Louis, MO, USA). Middlebrook 7H11 agar was purchased from BD Difco (NJ, USA). ADC growth supplement, Ziehl-Neelsen stain, rifampicin, and isoniazid were purchased from HiMedia (India). Calcium Chloride, Magnesium Sulphate, Sodium Chloride, Tween 80 and Glycerol were purchased from Fisher Scientific (MA, USA).

### 2.2 Bacterial cell culture and maintenance

Primary *Mycobacterium smegmatis* (mc^2^155) or *Mycobacterium tuberculosis* (H37Ra) cultures were grown in Middlebrook 7H9 broth supplemented with Glycerol, ADC and Tween-80 (0.1% v/v). A log phase primary culture was inoculated into a secondary culture without Tween-80 and supplemented with CaCl_2_ (2 mM) to promote an efficient infection of phages. Frozen stocks were prepared by adding 300 μL of a late log phase culture to 35% (v/v) final concentration of glycerol and stored at −80°C until future use. All cultures were maintained at 37°C with rotary shaking at 180 rpm. For optical density (OD) measurements, 300 μL of bacterial culture at various time intervals was diluted 10 times in media and pipetted several times to obtain a uniform cell suspension. Readings were taken using a spectrophotometer (Jenway 7205 UV/Visible Spectrophotometer) at 600 nm against a media blank.

### 2.3 Soft agar overlay and spot assays for phage enumeration

Soft agar overlay technique was used to detect and quantify phages. Briefly, soft agar was prepared by adding agar (0.8%) to 7H9 media along with the required supplements. The suspension was autoclaved and allowed to cool down to 42°C. 1 mL of late log phase bacterial culture (OD>1) and/or 100 μL of phage dilutions was added to 5 mL of the soft agar and was poured onto a 7H10 plate supplemented with CaCl_2_. The soft agar was allowed to cool down and solidify. The plate was incubated at 37°C overnight.

For verification of phage titers, spot assays were carried out in which late-log phase *M. smegmatis* cultures (OD>1) were plated on a Middlebrook 7H10 media plate supplemented with CaCl_2_ (2 mM) and glycerol to form a bacterial lawn. Phage samples were serially diluted and spotted on to the bacterial lawn. The plates were incubated at 37°C for 24 h and the number of plaques formed at various dilutions was counted.

### 2.4 Phage amplification and maintenance

Five phages (D29, TM4, Che7, PDRPv (Bajpai et al., 2018) and PDRPxv (Sinha et al., 2020)) used in this study were amplified on *M. smegmatis*. Similarly, three phages (D29, TM4 and DS6A) were amplified on *M. tuberculosis*. For liquid amplification, phage sample at 10 MOI was added to a log phase culture (with OD between 1-2) and incubated for 12-24 h at 37°C with rotary shaking at 180 rpm. The cultures were then centrifuged at 4000 g for 10 min and the supernatant was collected and filtered through a 0.22 μm syringe filter (Biofil, FPV203030-A). For plate amplification, a soft agar overlay containing log phase *M. smegmatis* (OD 1-2) was prepared and the phage sample (10 MOI) was spotted on top of the soft agar. The plates were incubated for 12-24 h at 37°C following which, the soft agar was collected into a 50 mL falcon tube. An appropriate amount of Magnesium Sulphate- Tris Chloride-Sodium Chloride Buffer (SM Buffer) was added to the falcon tube to completely immerse the soft agar and the falcon tube was incubated for 2-4 h at 37°C or overnight at 4°C. The falcon tube was then subjected to centrifuge at 4000 g for 10 min at 4°C and the supernatant was collected and filtered using 0.22 μm syringe filters. The soft agar overlay was used to determine the phage titers.

### 2.5 Bacterial growth kinetics

In order to study the bacterial growth kinetics, 7H9 broth supplemented with ADC and CaCl_2_ was prepared and inoculated with 50 μL of log-phase secondary bacterial culture (OD 1-2) per 10 mL of the media. The cultures were then incubated at 37°C with rotary shaking at 180 rpm. Periodic measurements of bacterial optical density (OD) were taken. For phage treated samples, a 10 MOI (for *M. smegmatis*) or 1 for (*M. tuberculosis*) of each phage in the respective phage solution was added at the appropriate time (either at the start of the experiment (lag phase) or at a specified interval of mid-log phase).

### 2.6 Bacterial growth kinetics with phage – pH, stationary phase and hypoxia

To study the effects of acidic pH on phage infection dynamics, phage kinetics assay was performed at varying the pH of the media. Briefly, the pH of the prepared media was reduced from 6.8 prior to the experiments using HCl to either a pH of 6.0 or pH of 5.5. The media was filtered and used for future growth kinetics experiments. Bacteria were inoculated into the media at pH 6.8, 6.0 and 5.5. For phage treated samples, a 10 MOI for each phage in the 5-phage cocktail was added at an appropriate time (either at the start of the experiment (lag phase) or at a specified interval of mid-log phase. OD was measured at 600 nm.

For stationary phase experiments, *M. smegmatis* cultures were grown in 7H9 media supplemented with Glycerol, ADC and CaCl_2_ (2 mM) until an OD of over 10 was reached. The stationary phase cultures were then either treated with the 5-phage cocktail at 10 MOI or left untreated. OD values were obtained at regular intervals at 600 nm.

For the studies involving hypoxia, the cultures were grown in plastic K2-EDTA Vacutainers (BD Cat No. 367525) to generate hypoxia. Briefly, K2-EDTA Vacutainers were obtained from BD and washed with autoclaved sterile water over a period of 24 h to remove the EDTA coating. The vacutainers were then washed with prepared media before experimentation. Upon inoculation, the vacutainers were sealed with the rubber septa caps and parafilm to prevent oxygen diffusion. Samples were collected through a syringe and 20G needle for spectrophotometer readings. Oxygen levels were monitored by adding Methylene Blue (0.01%) to the samples (Boshoff et al., 2004) and measuring the OD at 665 nm.

### 2.7 Ziehl-Neelsen staining

Ziehl-Neelsen staining was performed periodically to check for contamination and for the confirmation of *Mycobacterium spp*. as described by the manufacturer. Briefly, the bacterial suspension was spread evenly over the slide and air-dried for 30 min. The slide was then heat-fixed on a hot plate for 15 min at 60°C and then flooded with Carbol Fuchsin stain at 85°C. Care was taken not to overheat or boil the stain and the slide was frequently replenished with additional Carbol Fuchsin stain. The slide was allowed to remain on the hot plate for 10 min following which it was removed and allowed to incubate for an additional 5 min. The stain was washed away under running tap water and flooded with decolorizer for 2 min or until the smear was sufficiently decolorized. The slide was washed under running tap water and treated with Methylene Blue for 30 s. The slide was subjected to one final washing under running tap water to remove the Methylene Blue stain and was air-dried following which, it was observed under a 100× oil immersion objective using an upright fluorescence microscope (Olympus BX53F) for the presence of either pink rod-shaped acid-fast bacteria or blue stained contamination.

### 2.8 Antibiotic Minimum Inhibitory Concentration (MIC) determination

The MICs of both rifampicin and isoniazid were determined to establish the antimycobacterial activities of the respective antibiotics against *M. smegmatis* strain mc^2^ 155. Antibiotic stock solutions were prepared at concentration of 10 mg/mL in sterile deionized H_2_O for isoniazid and in DMSO for rifampicin. Stocks were stored at −20°C until further usage. Several dilutions were prepared in a 96- well plate for each of the antibiotic stock solutions to obtain various concentrations ranging from 1.5 μg/mL to 100 μg/mL. A total of 2×10^5^ cells of *M. smegmatis* were added to each of the wells. The 96-well plate was placed in a rotary shaker incubator at 37° C for 48 h and was observed for growth against control be measuring OD at 600 nm. The lowest concentration of antibiotic without bacterial growth was termed as MIC for the respective antibiotic.

### 2.9 Antibiotic growth kinetics

In order to study the effect of antibiotics-isoniazid and rifampicin on *M. smegmatis* growth, 7H9 broth supplemented with ADC and CaCl_2_ was prepared and inoculated with 50 μL of log-phase secondary bacterial culture per 10 mL of the media. The cultures were then incubated at 37°C with rotary shaking at 180 rpm. Periodic measurements of bacterial OD at 600 nm against a media blank. For antibiotic-treated samples, a final concentration of rifampicin at 40 μg/mL and 100 μg/mL and isoniazid at 40 μg/mL and 100 μg/mL were added at the appropriate time (either at lag phase or at a specified interval of mid-log phase). In order to study the potential synergistic relationship between antibiotics and phages, we co-treated bacterial cultures with rifampicin at 20 μg/mL and phage cocktail (at MOIs 0.01, 0.05 and 10) at either the lag phase or at a specified interval of mid-log phase. Periodic OD measurements were taken at 600 nm.

### 2.10 Phage kinetics against isoniazid resistant *M. smegmatis*

To study the effect of phage cocktail on antibiotic-resistant strain, 7H9 broth supplemented with ADC, CaCl_2_ and isoniazid (40 μg/mL) was prepared and inoculated with 50 μL of isoniazid-resistant (mc^2^ 4XR1) (Mohan et al., 2015; Padiadpu et al., 2015, 2016) log phase secondary bacterial culture per 10 mL of media. The cultures were then incubated at 37°C with rotary shaking at 180 rpm. Periodic measurements of bacterial OD were taken using a spectrophotometer at 600 nm against a media blank. For phage treated samples, the 5-phage cocktail was added at 10 MOI at an appropriate time (either at lag phase or at a specified interval of mid-log phase).

### 2.11 Checkerboard assay

To study the possible synergy between bacteriophages and rifampicin, checkerboard assays were performed. Briefly, *M. smegmatis* bacterial cultures were seeded into 48-well plates at a seeding density of 10^5^ colony forming units/mL. Respective concentrations of antibiotic and phages were added to the wells and the plates were incubated at 37°C with rotary shaking at 180 rpm for 48 h. OD readings were taken at 600 nm using a plate reader (Tecan Spark Multimode Microplate Reader).

### 2.12 Phage stability assay

To study the stability of the 5-phage cocktail at various pH, we incubated a 10^8^ solution at pH 6.8, 6.0 and 5.5 at 37°C. Samples were collected and titrated against *M. smegmatis* every 24 hours.

### 2.13 Statistical analysis

All experiments were performed on independent biological replicates. Statistical significance was determined for control and experimental groups using multiple t-test with Holm-Sidak method, with alpha = 0.05. For analysis of multiple groups, 2-way ANOVA was used with Sidak’s multiple comparisons test, with alpha = 0.05. GraphPad (Prism) was used for all statistical analysis. Data points were excluded if contamination was identified using acid-fast staining.

## 3 Results

### 3.1 Mycobacteriophages reduce bacterial growth in *M. smegmatis*

Several lytic Mycobacteriophages were procured and amplified to titers greater than 10^9^ pfu/mL. In order to test their efficacy, *M. smegmatis* bacterial cultures were infected with phages at the MOI of 10 either in the lag phase or in the mid-log phase. The optical density of the cultures at 600 nm was monitored using a spectrophotometer at various time intervals. Cultures treated with phages D29, TM4, Che7, PDRPv and PDRPxv individually showed a significant reduction in the optical density after 2-5 h when phages were added at log phase (time of addition shown by red arrows) (**Figure 1**). Similarly, when the phage infection was carried out at lag phase, no growth was observed for several hours. It is well known that bacteria evolve resistance to phages over time. *M. smegmatis* cultures infected with phages were also monitored for phage resistance. Phage resistance was observed in all phage treated *M. smegmatis* cultures. While D29 and Che7 treated cultures showed phage resistance at around 40 h, TM4 treated cultures showed phage resistance at 60 h; PDRPv and PDRPxv treated cultures showed phage resistance at 83 h and 101 h respectively (**Figure 1**). Bacterial cultures infected with phages at an MOI of 30 showed no significant increase in the onset of phage resistance compared to MOI of 10 (**Supplementary Figure S1**). To test if the bacteria growing after the phage challenge had any altered susceptibility to the phages, we titrated our individual phage stocks on bacterial growths obtained after treatment with D29 and PDRPv. While D29 and PDRPv showed much reduced titers (2-3 log orders) against their respective regrowing bacteria, no significant reduction in susceptibility to other phages was observed (**Supplementary Figure S2**).

**Figure 1.**
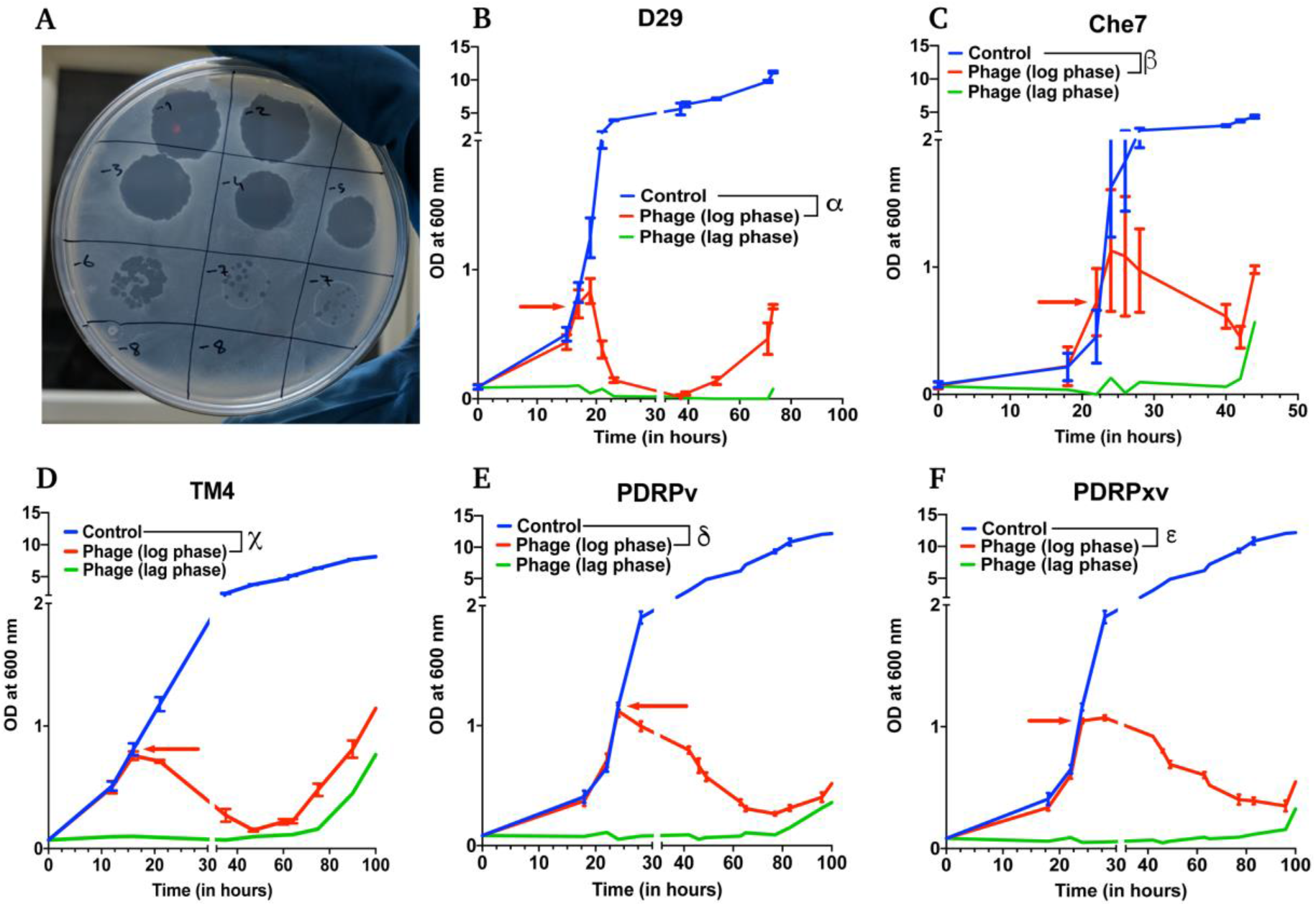
Efficacy of infection and evolution of phage resistance is dependent on individual Mycobacteriophages. **A)** Photograph showing spot clearance of a lawn of *M. smegmatis* after treatment with D29. Growth curves of *M. smegmatis* in liquid culture in the presence of **B)** D29, **C)** Che7, **D)** TM4, **E)** PDRPv and **F)** PDRPxv phage. Red arrow represents the time point at which phage solution was added to the cultures. These experiments were repeated several times (≥2) and the representative graphs were plotted. Multiple t-test with Holm-Sidak method was used to estimate the statistical significance. **α** represents a significant difference (p<0.05) between control and D29 treated (log phase) for all time points between 21 h and 73 h. **β** represents a significant difference (p<0.05) between control and Che7 treated (log phase) for all time points between 28 h and 44 h. **χ** represents a significant difference (p<0.05) between control and TM4 treated (log phase) for all time points between 21 h and 105 h. **δ** represents a significant difference (p<0.05) between control and PDRPv treated (log phase) for all time points between 28 h and 106 h. **ε** represents a significant difference (p<0.05) between control and PDRPxv treated (log phase) for all time points between 24 h and 106 h.

### 3.2 Phage cocktails show synergy in growth reduction

To test the effect of phage cocktails, *M. smegmatis* cultures infected with multiple phages (a cocktail of 3 or a cocktail of 5) at 10 MOI of each phage were monitored for OD at 600 nm (**Figure 2**). Phage cocktail treatment significantly reduced the optical density of the bacterial cultures compared to controls 5 h post treatment. Phage resistance in cocktail treated cultures prolonged to 105 h in case of 3 phage cocktail (D29, TM4 and Che7) (**Figure 2A**) and to 322 h in case of 5 phage cocktail (D29, TM4, Che7, PDRPv and PDRPxv) (**Figure 2B)**. The corresponding Colony Forming Units (CFU)/mL readings were obtained and the viable cell count was observed to be correlating with OD readings (**Supplementary Figure S3A**). This shows that the cocktail of phages can act synergistically to delay the evolution of resistance. Periodic phage titer measurements showed increase in the phage titers as the bacterial OD dropped and phages could be detected upto 400 h although a steady decline in the titer was observed (**Supplementary Figure S7A**). Subsequent phage stock rechallenge on the growing bacteria in the 5-phage cocktail kinetics showed that all phage stock titers had dropped by >99% (2-3 log order) on the pre-exposed bacteria compared to untreated bacteria **(Supplementary Figure S3B)** suggesting evolution of the bacteria to resist phage mediated lysis. It is important to note that the phages used here belong to different genomic clusters: cluster A (D29, subcluster: A2), cluster K (TM4, subcluster: K2) and cluster B (PDRPv and PDRPxv, both in subcluster B1) (2–6). Cluster information for Che7 is not known. Hence, it is expected that these phages will employ different mechanisms to infect the bacteria which is consistent with delay in evolution of resistance (**Figure 2**) as more phages are added to the cocktail. The 5-phage cocktail was selected and used for future experiments. To determine the time it takes for OD of bacterial cultures to start reducing after phage addition, we took OD readings every 15-20 mins. We found that OD of the bacterial cultures starts to drop only after 120-150 mins **(Supplementary Figure S4)** which is consistent with the previously reported latency period of D29 (~60 min) (Samaddar et al., 2015; Bavda and Jain, 2020) and PDRPxv (~135 min) in *M smegmatis* (Sinha et al., 2020).

**Figure 2.**
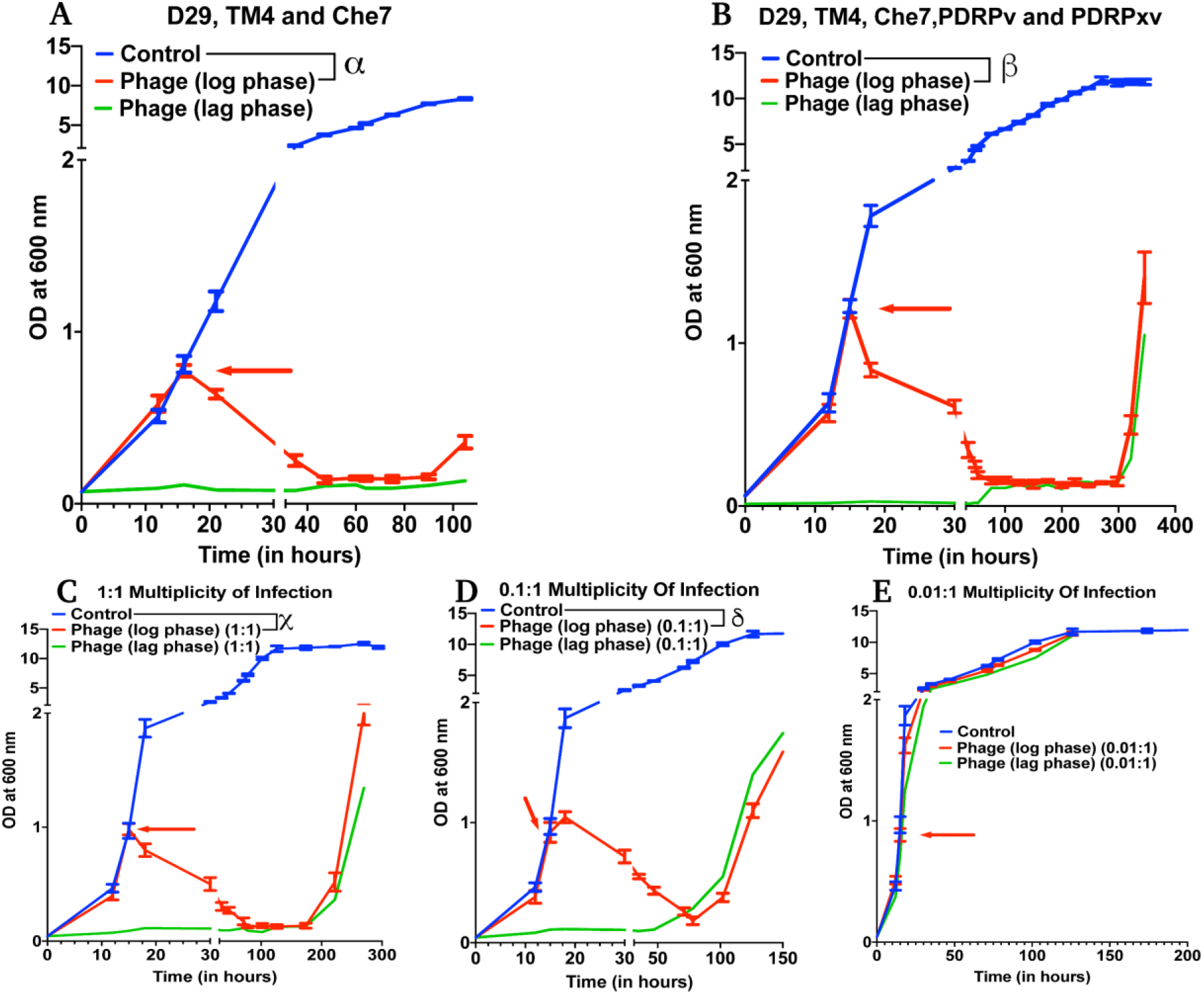
Phage cocktails show synergy compared to individual phage and their efficacy is MOI dependent. Growth kinetics of *M. smegmatis* cultures treated with **A)** 3-phage cocktail (D29, Che7 and TM4) and **B)** 5-phage cocktail (D29, TM4, Che7, PDRPv and PDRPxv) at MOI of 10 for each phage against a non-treated control. Growth kinetics of *M. smegmatis* cultures treated with a 5-phage cocktail at the MOI of **C)** 1, **D)** 0.1 and **E)** 0.01 for each phage in the phage cocktail. The experiments with MOI of 10 with 3- and 5-phage cocktails were repeated twice and the representative graphs are shown here. Red arrow represents the time point phage solution was added to the cultures. Multiple t-test with Holm-Sidak method was used to estimate the statistical significance (p<0.05). **α** represents a significant difference (p<0.05) between control and three-phage cocktail (D29, TM4 and Che7) treated (log phase) for all time points between 21 h and 105 h. **β** represents a significant difference (p<0.05) between control and five-phage cocktail (D29, TM4, Che7, PDRPv and PDRPxv) treated (log phase) for all time points between 18 h and 346 h. **χ** represents a significant difference (p<0.05) between control and 5 phage cocktail treated (MOI 1, log phase) for all time points between 18 h and 270 h. **δ** represents a significant difference (p<0.05) between control and 5 phage cocktail treated (MOI 0.1, log phase) for all time points between 18 h and 174 h.

To test for the efficacy of mycobacteriophages at various Multiplicity of Infection (MOI), *M. smegmatis* cultures were infected with a MOI of 10, 1, 0.1 and 0.01 of each phage: bacteria in either the lag phase or the mid-log phase. A 10 MOI **(Figure 2B)** significantly reduced the growth of bacterial culture 3 h post treatment with phage resistance emerging at 322 h. A 1 MOI **(Figure 2C)** also resulted in a significant reduction of bacterial growth 3 h post treatment, however, phage resistance emerged at around 222 h. While a 0.1 multiplicity of infection **(Figure 2D)** resulted in a reduction in growth, resistance was observed at 102 h for mid-log phase treated cultures. A 0.01 MOI did not result in any significant reduction in the bacterial growth **(Figure 2E)**. For all future experiments, we have used the MOI of 10 to evaluate the maximum lysing potential of phages against Mycobacterium.

### 3.3 Mycobacteriophages are effective in under low pH and hypoxia

Several species of Mycobacterium exist as intracellular pathogens in acidic and hypoxic environments (Sibling et al., 1987; Oh and Straubinger, 1996; Rastogi et al., 2001; Houben et al., 2006; Benítez-Guzmán et al., 2019; Seto et al., 2020). In order for phage therapy to be efficient against such mycobacterial infections, phages need to retain their activity and infectivity in these extreme environments. We, therefore, evaluated the efficiency of our 5-phage cocktail under low pH conditions normally observed in infected phagosomes (ranges from pH 4-6.5) of mammalian cells. *M. smegmatis* cultures did not grow at pH 5.0 and below. Hence, we chose pH 5.5 and 6.0 to test the effectiveness of phages as these pH ranges are known to exist in early and late endosomes (Rohde et al., 2007). We tested the phage stability in 7H9 medium with pH 6.8, 6.0 and 5.5. Although there was a progressive loss of phage titers over 300 h, no significant difference in phage activity was observed at lower pH compared to pH of 6.8 **(Supplementary Figure S5)**. Under low pH conditions of 6.0 and 5.5, the mycobacteriophages remain efficient in infecting and killing *M. smegmatis* and showed resistance around 300 h, similar to those observed at pH 6.8 (standard pH of 7H9 medium) (**Figure 3A-B, Supplementary Figure S6A-B)**.

**Figure 3.**
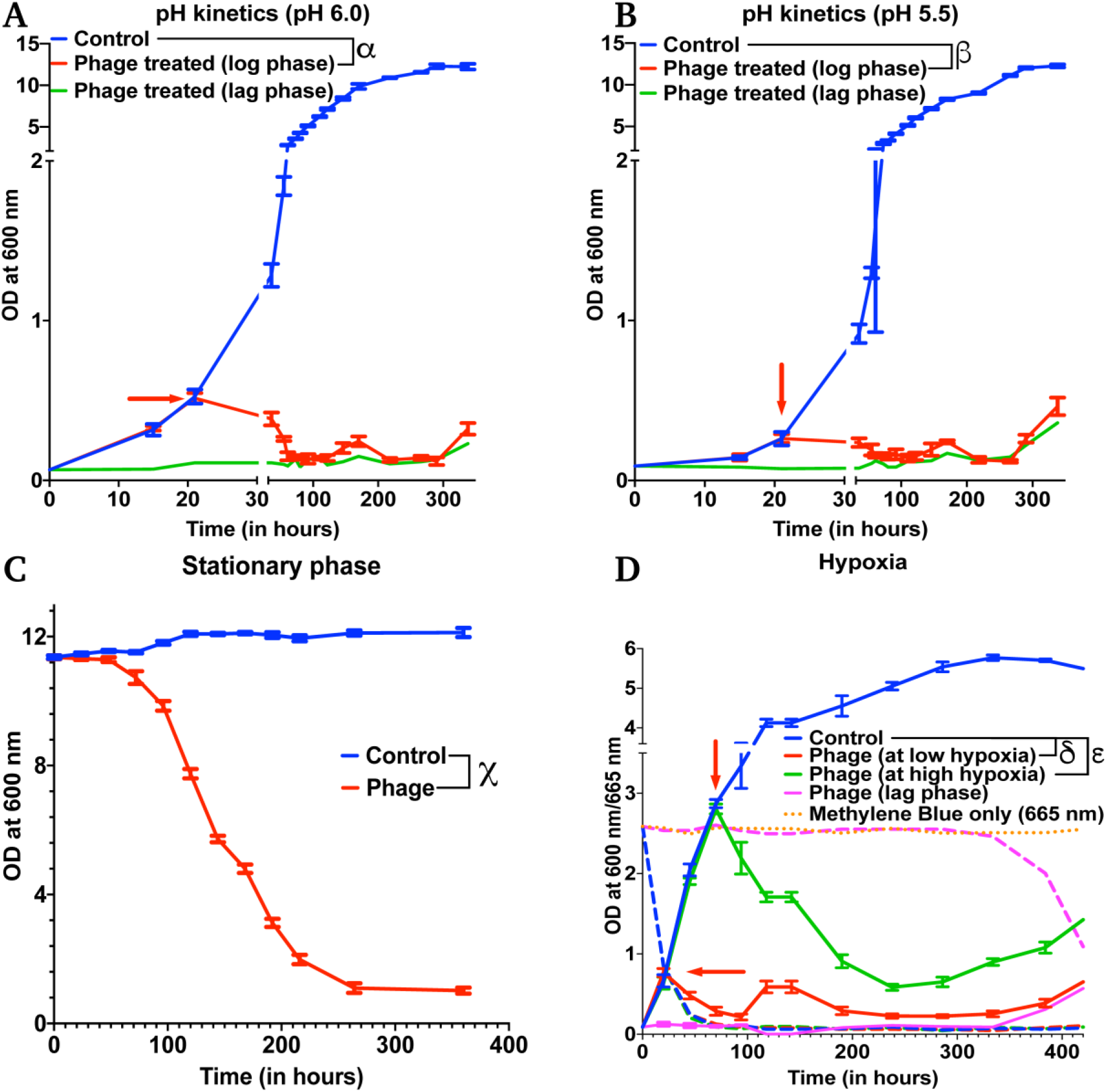
Mycobacteriophages are effective under various pathophysiological conditions. Growth kinetics of *M. smegmatis* cultures treated with a 5-phage cocktail at **A)** pH condition of 6.0, **B)** pH condition of 5.5, **C)** stationary phase and **D)** hypoxic conditions. For hypoxia experiment, solid lines represent bacterial OD at 600 nm and dashed lines are the respective Methylene Blue readings at 665 nm. Red arrow represents the time point phage solution was added to the cultures. These experiments were repeated twice and the representative graphs are shown here. For **A, B** and **C**, multiple t-test with Holm-Sidak method was used to estimate the statistical significance (p<0.05). **α** represents a significant difference (p<0.05) between control and phage cocktail treated (pH 6.0, log phase) for all time points between 36 h and 338 h. **β** represents a significant difference (p<0.05) between control and phage cocktail treated (pH 5.5, log phase) for all time points between 73 h and 338 h. **χ** represents a significant difference (p<0.05) between control and phage cocktail treated (stationary phase) for all time points between 24 h and 360 h. For **D**, 2-way ANOVA with Sidak’s multiple comparisons test was used to determine statistical significance (p<0.05). **δ** represents a significant difference between control and phage (at low hypoxia) for all time points between 45 h and 432 h, and **ε** represents a significant difference between control and phage (at high hypoxia) for all time points between 94 h and 432 h.

Efficacy of the phages under hypoxic conditions was checked by culturing bacteria in sealed vacutainers with septa. Since, there is no exchange of gases in this model, as the bacteria grows over time, the oxygen present in the tube depletes creating a hypoxic environment. This was confirmed by adding methylene blue to the media that turns colorless in the absence of oxygen (**Supplementary Figure S8)** (Sumitani et al., 2004). We infected cultures with the 5-phage cocktail, at either low hypoxia (intermediate methylene blue absorbance) or high hypoxia (at lowest methylene blue absorbance). We observed efficient lysis of bacteria by the 5-phage cocktail in both the hypoxic conditions 20-25 h post treatment. (**Figure 3D** and **Supplementary Figure S6D**). Similar to pH experiments, the emergence of phage resistance was observed in both low and high hypoxia phage treated samples in ~300 h.

### 3.4 Mycobacteriophages are effective in stationary phase of bacterial growth

Within the host environment, Mycobacterial species slow their growth rate and often enter a non-replicative state (Voskuil et al., 2004; Houben et al., 2006; Ehlers and Schaible, 2012). This reduced growth state poses a significant hurdle in the treatment of infections with conventional antibiotics (Gutierrez et al., 2017). For phage therapy to be efficient against mycobacterial infections, phages need to retain the infectivity against the slow growing bacteria. To test the effect of phage cocktail against *M. smegmatis* in stationary phase, *M. smegmatis* cultures were allowed to reach an OD>10. The stationary phase cultures were then infected with the 5-phage cocktail without any supplementation with nutrients and periodic OD measurements were taken at 600 nm. Significant reduction in OD and CFU were observed in phage treated samples 24 h post treatment, demonstrating that the 5-phage cocktail is effective in stationary phase conditions (**Figure 3C, Supplementary Figure S6C**). Interestingly, even after 350 h of culture, no growth of bacteria was observed. In periodic phage titer measurements, phages could be detected upto 400 h although a steady decline in the titer was observed **(Supplementary Figure S7B)**.

### 3.5 Efficacy of isoniazid and rifampicin against *M. smegmatis*

Tuberculosis infections are conventionally treated by a combination of antibiotics, including the first-line drugs, isoniazid and rifampicin (Geneva: World Health Organisation, 2019). To compare our phage kinetics with these antibiotics, we first determined the minimal inhibitory concentration of rifampicin (MIC: 25 μg/mL) and isoniazid (MIC: 15 μg/mL) on *M. smegmatis* (mc^2^155). We then treated the cultures with 40 μg/mL and 100 μg/mL of the respective antibiotic at either lag phase or at mid-log phase. For 40 μg/mL antibiotic concentration, there was no significant reduction of bacterial growth at log phase (**Supplementary Figure S9 A, B**). For 100 μg/mL, we observed only a transient growth inhibition of *M. smegmatis* compared to the phage treatment (**Supplementary Figure S9 C, D**). The cultures continue to grow even in the presence of high concentration (several times the MIC values) of antibiotics. This growth inhibition was much more pronounced in lag-phase cultures treated with antibiotics than log-phase cultures treated with antibiotics. Antimicrobial susceptibility depends on multiple factors such as inoculum size, metabolic state of the bacteria and cellular respiration rate (Lobritz et al., 2015; Li et al., 2017; Lopatkin et al., 2019). All these factors change as bacterial cells transition from lag phase to exponential growth and antibiotic efficacy may be variable even at the same concentration for different growth states and bacterial numbers. Isoniazid interferes with mycolic acid synthesis and hence is more effective on replicating Mycobacteria but due to high numbers of bacteria in the exponential phase, its efficacy was lower than in the lag phase. These results are also consistent with the previously observed results that isoniazid had reduced efficacy on exponentially growing *M. smegmatis* cultures (Teng and Dick, 2003).

### 3.6 Testing synergy between Mycobacteriophages and antibiotics

To check whether treatment with the 5-phage cocktail enhances the efficacy of controlling bacterial growth in antibiotic-treated cultures, we added 20 μg/mL of rifampicin (~MIC value) along with a 5-phage cocktail solution at an MOI of 0.05, 0.01 or 10 during the lag phase or the mid-log phase. For MOI of 0.01, we did not observe a significant reduction in growth in only phage treated samples at both lag and log phases compared to untreated samples. However, rifampicin and phage cotreated samples showed a significant growth reduction in the log phase 6 h post treatment with resistance developing around 224 h **(Figure 4A)**. In the lag-phase bacteria, phage addition at 0.01 MOI had no effect on growth and rifampicin treated bacteria showed growth within 50 h while the lag-phase samples co-treated with both rifampicin and phage cocktail showed no growth emergence upto 254 h. This prominent reduction in bacterial growth in cultures co-treated with rifampicin and phage cocktail could be attributed to antibiotic-phage synergy previously observed in various other bacterial infections including *P. aeruginosa* infection treatment (Oechslin et al., 2017), *S. aureus* infection treatment (Chhibber et al., 2013) and *A. baumannii* infection treatment (Jansen et al., 2018). The effectiveness of a cotreatment at low MOI of phage cocktails and rifampicin concentrations shows phage-antibiotic synergy. While rifampicin binds to RNA polymerase that might interfere with phage amplification, a staggered treatment with phages followed by rifampicin has been shown to be synergistic (Rahman et al., 2011; Tkhilaishvili et al., 2018). Similar observations have been noted for antibiotics that target the DNA replication and translational machinery (Torres-Barceló et al., 2014; Kim et al., 2018). It is possible that the co-incubation of both anti-bacterial agents prevents the emergence of bacteria that is tolerant to the individual anti-bacterials and hence show synergy. For cultures treated with a phage cocktail MOI of 0.05, we observed a similar reduction in growth with phage resistance emerging at 254 h in log-phase treated samples (**Supplementary Figure S10A**). Interestingly, the phage-antibiotic synergy at log-phase became less pronounced with increase in phage MOI. At MOI of 10, no synergy was observed in log phase cultures but a delay in emergence of resistance in lag phase treated cultures was observed **(Supplementary Figure S10B).** To further test the phage synergy with rifampicin, we performed experiments at different concentration of phage-rifampicin through a checkerboard assay (**Figure 4B**) and found their interaction to be synergistic in preventing bacterial growth (FIC index of 0.43 (<0.5) at phage MOI 0.1 and rifampicin concentration of 10 μg/mL) (Hall et al., 1983).

**Figure 4.**
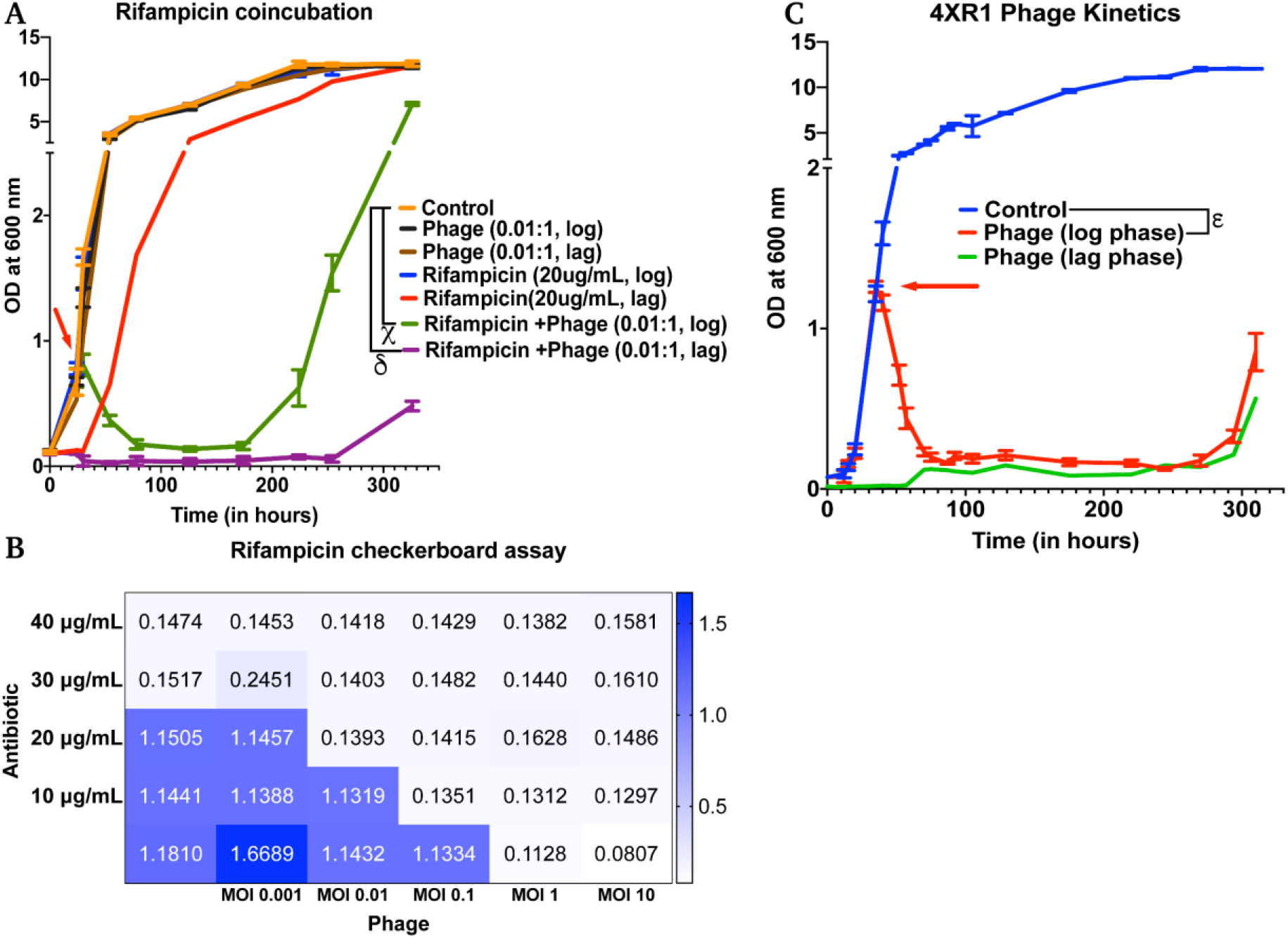
Phage cocktails show synergy with antibiotics and are effective against antibiotic-resistant Mycobacterium. **A)** 5-phage cocktail (MOI 0.01) and rifampicin (20 μg/mL)- **χ** and **δ** represent a significant difference between control and rifampicin + phage (log phase), and rifampicin + phage (lag phase) for all time points between 30 h and 326 h. **B)** Checkerboard assay of M. smegmatis cultures treated with rifampicin and 5-phage cocktail (D29, TM4, Che7, PDRPv and PDRPxv). **C)** Growth kinetics of isoniazid-resistant M. smegmatis (4XR1) cultures treated with a 5-phage cocktail. represents a significant difference between control and phage treated (log phase) for all time points between 40 h and 310 h. Red arrow represents the time point antibiotic/phage solution was added to the cultures. For A, 2-way ANOVA with Sidak’s multiple comparisons test was used to determine statistical significance (p<0.05). For C, multiple t-test with Holm-Sidak method was used to estimate the statistical significance (p<0.05).

### 3.7 Mycobacteriophages are able to lyse antibiotic-resistant *M. smegmatis*

Isoniazid-resistant infections are observed to be on the rise across the globe (Geneva: World Health Organisation, 2019). We, therefore, looked at the efficacy of phage infection and mycobacteriophages against isoniazid-resistant strain of *M. smegmatis* (Mohan et al., 2015; Padiadpu et al., 2015, 2016). First, we tested whether individual phages have altered efficiency in infecting and lysing the isoniazid-resistant strain. The phages retained their initial high titers against isoniazid-resistant *M. smegmatis* and no significant drop in the efficiency of plating was observed **(Supplementary Figure S11)**. Next, we observed that the 5-phage cocktail remains effective in infecting and lysing antibiotic-resistant *M. smegmatis* (**Figure 4C**). We observed a sharp decrease in the OD of phage-treated log-phase cultures 5 h post treatment and an extended delay in the growth of lag-phase cultures. We also observed growth of phage treated bacterial cultures at 294 h, owing to the emergence of phage resistance similar to the previous experiments.

### 3.8 Mycobacteriophages are effective against *M. tuberculosis*

Next, we checked whether other slow-growing species of Mycobacteria also exhibit similar dynamics with Mycobacteriophages. Although several reports have shown that phages infect and kill *M. tuberculosis,* however, whether cocktail of phages can prevent growth of *M. tuberculosis* strains over long durations and how long it takes for evolution of resistance is not known. In our 5-phage cocktail, two phages (D29 and TM4) are known to infect *M. tuberculosis* strains. These two phages and another phage DS6A were amplified using *M. tuberculosis* (H37Ra) as a host because the phages expanded on *M. smegmatis* had low titer on H37Ra. For infection experiments, a phage cocktail was prepared using these three phages. Since our phage titers against *M. tuberculosis* strains were low (~10^7^ PFU/mL) even after several rounds of amplification, we used a low inoculation density (~10^4^ CFU/mL) of bacteria to perform infection experiments at 10 MOI for each phage. We proceeded to treat the cultures of *M. tuberculosis* (H37Ra strain) with this three-phage cocktail in both the lag phase (OD ~0.06) and in the log phase. Phage treatment was successful and efficient in preventing the growth of *M. tuberculosis* for several weeks (**Figure 5A)**. Emergence of phage resistance was observed after 1368 h (57 days). We also observed reduction in bacterial CFU after phage treatment, consistent with our OD measurements **(Figure 5B**). These experiments show that phages can be also effective against slow growing Mycobacterium for long durations.

**Figure 5.**
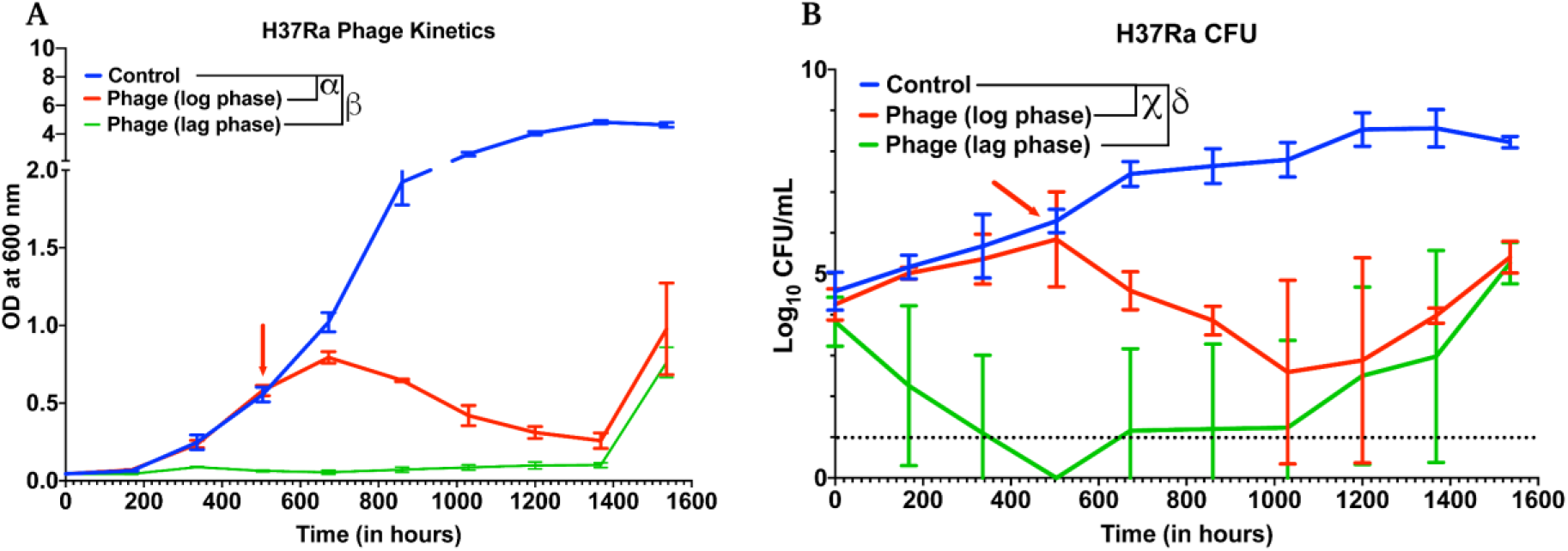
Phage cocktails are effective against M. tuberculosis. **A)** Growth kinetics of *M. tuberculosis* (H37Ra) after treatment with a 3-phage cocktail (D29, Che7 and DS6A). Multiple t-test with Holm-Sidak method was used to estimate the statistical significance (p<0.05). **α** represents a significant difference between control and phage treated group (log phase) for all time points between 672 h and 1536 h. **β** represents a significant difference (p<0.05) between control and phage treated group (lag phase) for all time points between 168 h and 1536 h. **B)** Colony Forming Unit (CFU) measurements over time of *M. tuberculosis* cultures treated with a *3*-phage cocktail. **χ** represents a significant difference (p<0.05) between control and phage (log phase) for all time points between 672 h and 1536 h. **δ** represents a significant difference (p<0.05) between control and phage (lag phase) for all time points between 168 h and 1536 h. Multiple t-test with Holm-Sidak method was used to determine the statistical significance. The detection limit for the assay is represented in black dotted line (10 CFU).

## 4 Discussion

Phage therapy has been successfully used in the treatment of various bacterial infections including *P. aeruginosa* urinary tract infection (Khawaldeh et al., 2011), multidrug-resistant *A. baumannii* infection (Schooley et al., 2017), *P. aeruginosa* aortic graft infection (Chan et al., 2018), and more recently, *Mycobacterium abscessus* infection of a 15- year old cystic fibrosis patient (Dedrick et al., 2019). Although phage therapy holds strong potential, there is little understanding of phage-host dynamics in disease environments. In this study, we studied the host-phage infection dynamics under various physiological and disease-mimicking conditions.

We chose mycobacterial species as our phage host, as persistent and multidrug-resistant mycobacterial infections have been increasing in incidence over the past decades and there is a need for new therapy against such infections (Iseman, 1994; Gygli et al., 2017; Aslam et al., 2018; Cambau et al., 2018; Luthra et al., 2018). For phage therapy to be successful and efficient against mycobacterial infections, the phages need to retain their infectious activity in these extreme environments and infect non-replicative stationary phase bacteria. D29 is a well-known phage against *M. tuberculosis* known to produce superoxide radicals in oxygen-rich environments, which synergizes with phage mediated lysis to accelerate the killing of Mycobacterium (Samaddar et al., 2016). However, it is observed to be ineffective in hypoxic conditions (Swift et al., 2014). TM4, on the other hand, is shown to be efficient in infecting *M. tuberculosis* during stationary phases and hypoxic conditions (Swift et al., 2014). Biochemical analysis revealed that a peptidoglycan motif in the tape measure protein of TM4 phage confers an advantage in infecting bacteria in stationary phase (Piuri and Hatfull, 2006). This complementary nature of phages suggests that a cocktail of phages would be effective in the elimination of mycobacterial infections across a broad range of environments and growth states. Although research has been carried out on individual mycobacteriophages under specific environments, limited information exists which solidifies our understanding of phage cocktail infection kinetics and evolution of phage resistance under different disease-mimicking conditions.

In this study, we show that a 5-phage cocktail is efficient against *M. smegmatis* at various pH, low oxygen availability and different growth rate of bacteria. Furthermore, for the first time, our study shows the *M. tuberculosis* growth kinetics in presence of phage cocktail over long durations. We found that a 3-phage cocktail of D29, TM4 and DS6A was successful in preventing as well as reducing the growth of *M. tuberculosis* (H37Ra) for several weeks (**Figure 5**).

We also found that two frontline TB antibiotics-isoniazid and rifampicin were limited in their ability to control the growth of *M. smegmatis* in the exponential phase of bacterial growth (**Supplementary Figure S9**). The more striking result from our data was the synergy of phages with rifampicin and effectiveness of phages on isoniazid-resistant *M. smegmatis* (**Figure 4A-C**). This finding stands in line with observations of synergy during phage-antibiotic combination treatments for other bacterial systems (Comeau et al., 2007; Chhibber et al., 2013; Oechslin et al., 2017; Jansen et al., 2018). Although phage therapy holds promise in treating bacterial infections and antibiotic resistance, bacteria are known to develop resistance against phages through various mechanisms (Labrie et al., 2010). However, unlike antibiotics which have limited classes and require scrutinized drug design, phages are abundant in nature (Edwards and Rohwer, 2005) and the mycobacteriophage repository alone is over 10,000 in number (Russell and Hatfull, 2017), providing a much larger repertoire of arsenal against bacterial infections. However, when designing a phage cocktail, the phage resistance needs to be considered. The phage-bacterial dynamics and evolution of resistance described here are some the first studies performed on mycobacterial species and provide useful information for future exploration of phages for therapeutic use. We found that the bacteria eventually evolve resistance to the phage or phage cocktail as is seen by regrowth of the bacteria after phage challenge even though the phages can be detected in the medium at all times. Subsequent challenge of our phage stocks on this regrowing population showed a drop of 2-3 log orders (>99%) in titers suggesting evolution of the bacteria to resist phage mediated lysis. The mechanisms of this resistance are not clear and need further investigation. Bacterial hosts are known to develop resistance against phages through various mechanisms including genetic mutations, hypermutable loci, the blockage of the injection of phage DNA, cleavage of injected phage DNA, interference with phage assembly, abortive infection and other phage defense systems such as the R-M system, DISARM system or the CRISPR-Cas system (Azam and Tanji, 2019; Turkington et al., 2019). Whether *Mycobacterium spp.* utilize these mechanisms to overcome the phage challenge is not clear and requires further research. In our studies, we observed a strong correlation between the delay in the emergence of phage resistance and the number of phages used in our phage cocktail.

Our phage infection at different MOI suggests that it is important to deliver a high number of phages at the site of infection for therapy to be successful (**Figure 2C-D**). Several approaches have been developed to deliver active phages to the deep lung such as vibrating mesh nebulizer, soft mist inhalers and dry powder inhalers (Puapermpoonsiri et al., 2009; Vandenheuvel et al., 2013; Liu et al., 2016; Carrigy et al., 2017; Agarwal et al., 2018). Similarly, delivery to macrophages by direct phagocytosis of phages (Jończyk-Matysiak et al., 2015) or through nanoparticles/microparticles can result in targeting of the intracellular niche of the bacteria (Fenaroli et al., 2014; Nieth et al., 2015; Agarwal et al., 2018). However, the current understanding of the diffusion of phages across the granuloma remains limited and merits further studies in animal models.

A limitation to this study is that most experiments were performed on non-pathogenic, fast growing *M. smegmatis* and may not translate to slow growing pathogenic species such as *M. tuberculosis*. Although we show that phages can infect and lyse *M. tuberculosis* for long durations, further studies are warranted in testing the efficiency of these phages on such slow growing pathogenic species for longer durations and in acidic and hypoxic conditions. However, efficacy of mycobacteriophages in lysing intracellular slow growing mycobacteria *in vitro* as well as *in vivo* has been demonstrated by other groups. Preliminary reports on D29 incubation with H37Rv infected mouse macrophages show reduction in intracellular bacterial burden, however penetration of D29 inside the macrophage was reported to be low (Lapenkova et al., 2018). It is critical for the phages to directly encounter bacteria to be effective. Phage delivery inside infected mammalian cells can be facilitated by using carriers, for instance by adsorbing on non-pathogenic bacteria. This led to enhanced intracellular localization of phages *in vitro* in *M. avium* and *M. tuberculosis* infected macrophages and reduction in bacterial load in a mouse model of disseminated *M. avium* infection (Broxmeyer et al., 2002; Danelishvili et al., 2006). Bactericidal efficiency of phages has also been shown *in vivo* in a *M. ulcerans* infected mouse footpad model (Trigo et al., 2013). Another challenge for phage therapy is the human immunological response. Although phages are known to elicit immune response (Ooi et al., 2019), previous studies have shown that phages and host immune response can synergistically result in the elimination of acute and chronic bacterial infections (Leung and Weitz, 2017; Roach et al., 2017; Waters et al., 2017). This is further supported by the successful treatment of several patients after phage administration in clinics.

## Conclusion

In summary, we demonstrate the efficacy of phages under various pH ranges (6.8, 6.0 and 5.5), stationary phase as well as under hypoxic conditions. We show that phages are effective in infecting and lysing antibiotic resistant bacteria and show synergy with antibiotics. We further show that a three-phage cocktail is effective in preventing the growth of slow growing *M. tuberculosis* for several weeks.

## Author Contribution

RA conceived the idea. YCK and RA designed the research. YCK and PRS performed the research. YCK and RA analysed the data and wrote the manuscript.

## Acknowledgements

We acknowledge Dr Sujoy K. Das (Bose Institute, Kolkata), Dr Urmi Bajpai (Acharya Narendra Dev College, Delhi), Dr V.N. Azger Dusthackeer (National Institute for Research in Tuberculosis, Chennai) and Dr Graham Hatfull (University of Pittsburgh) for providing us with mycobacteriophages. We are grateful towards Dr Amit Singh (Indian Institute of Science, Bengaluru) for providing bacterial strains. We thank Dr Nagasuma Chandra for kindly providing us with the isoniazid-resistant *M. smegmatis*. We thank the Molecular Reproduction, Development and Genetics, and Materials Engineering common facility for instrument access.

