## Supplementary Information for "Antimycobacterial potential of Mycobacteriophage under disease-mimicking conditions"

**Running Title: Antimycobacterial potential of phage**

### Supplementary Information

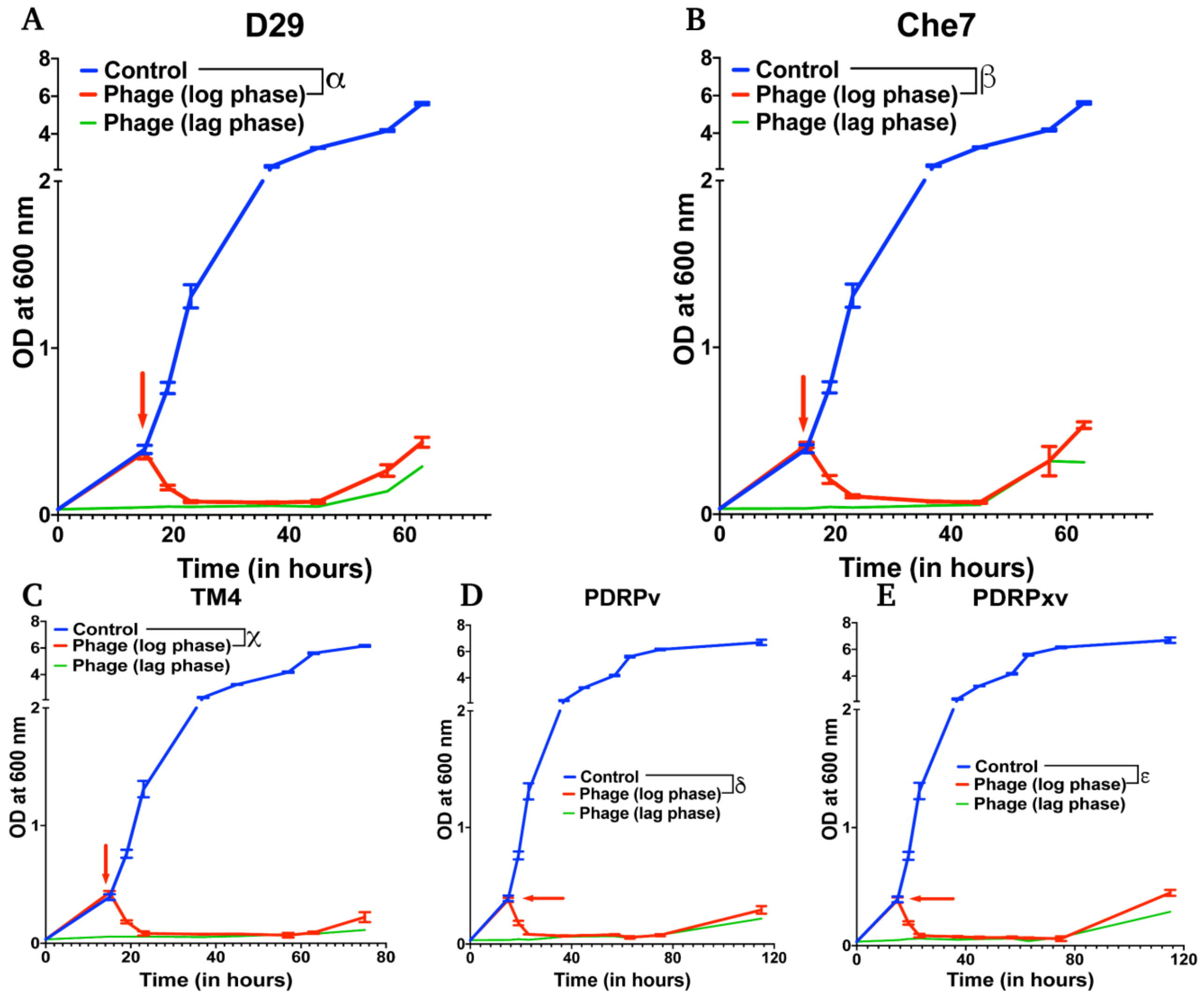


**Supplementary figure S1. Growth kinetics of M *smegmatis* at MOI 30** of A) D29 B) Che7 C) TM4 D) PDRPv and E) PDRPxv**.** Red arrow represents the time point at which phage solution was added to the cultures. Multiple t-test with Holm-Sidak method was used to estimate the statistical significance. $\alpha$ represents a significant difference ($p<0.05$) between control and D29 treated (log phase) for all time points between 19 h and 63 h. $\beta$ represents a significant difference ($p<0.05$) between control and Che7 treated (log phase) for all time points between 19 h and 63 h. $\chi$ represents a significant difference ($p<0.05$) between control and TM4 treated (log phase) for all time points between 19 h and 75 h. $\delta$ represents a significant difference ($p<0.05$) between control and PDRPv treated (log phase) for all time points between 19 h and 115 h. $\varepsilon$ represents a significant difference ($p<0.05$) between control and PDRPxv treated (log phase) for all time points between 19 h and 115 h.

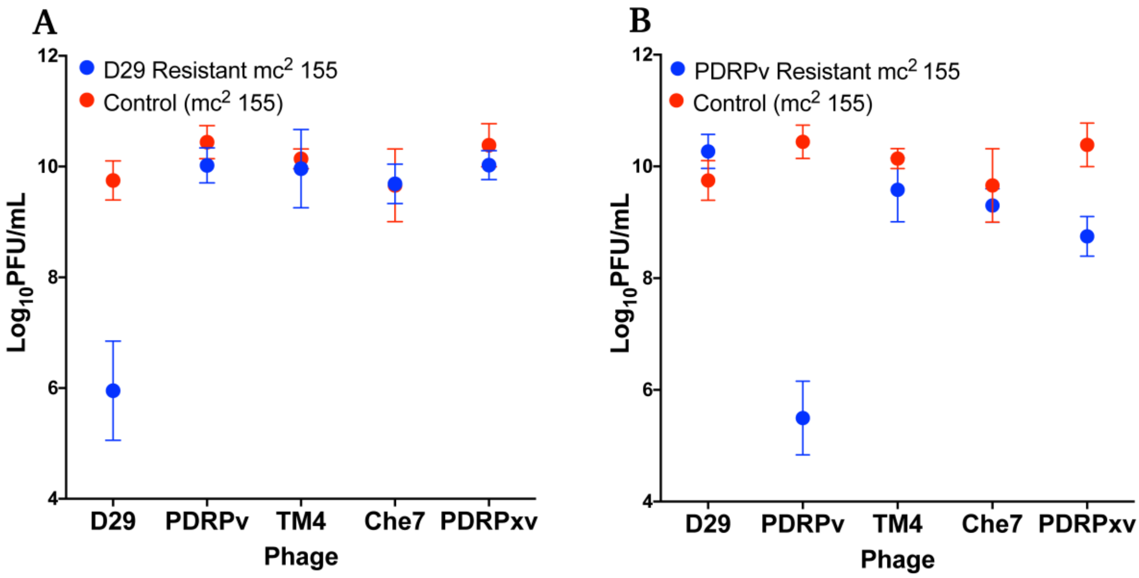


**Supplementary figure S2. Phage exposure causes reduction in respective phage titer on regrowing bacteria.** Phage susceptibility of all phages on **A)** D29 exposed *M. smegmatis* and **B)** PDRPv exposed *M. smegmatis*. Data are plotted as mean ± SD.

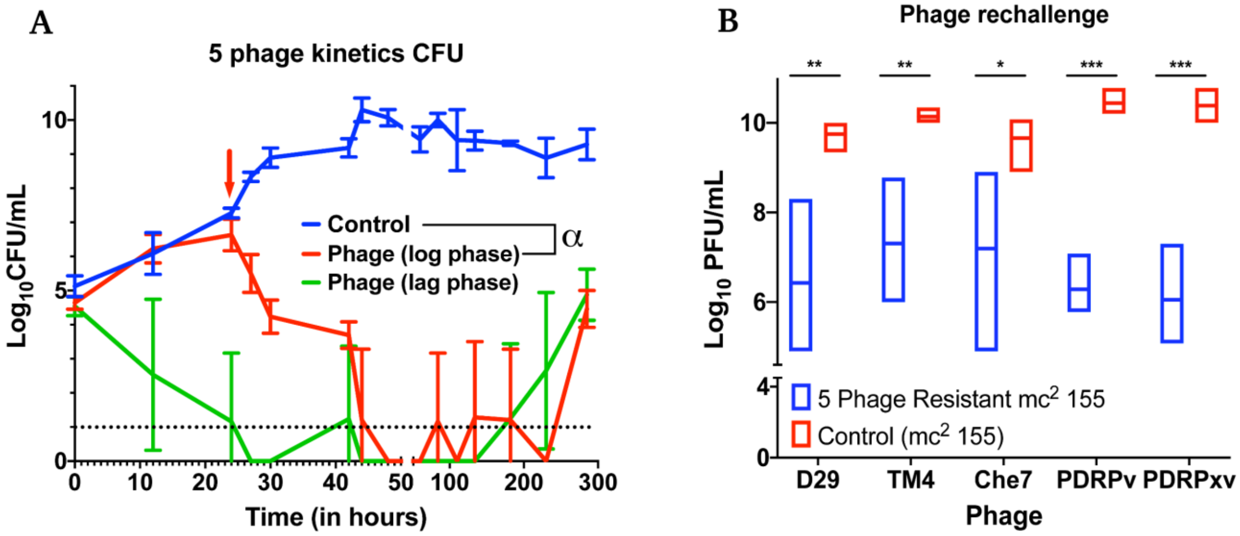


**Supplementary figure S3. Phage cocktail show synergy compared to individual phage and its efficacy is MOI dependent. A)** Colony Forming Unit (CFU) measurements of *M. smegmatis* over time for the 5-phage cocktail (D29, TM4, Che7, PDRPv and PDRPxv) against a non-treated control. α represents a significant difference ($p<0.05$) between control and phage (log phase) for all time points between 27 h and 285 h. Red arrow represents the time point phage solution was added to the cultures. **B)**

Individual phage titers against the phage resistant growth observed in the 5-phage kinetics. Multiple t-test with Holm-Sidak method was used to determine the statistical significance (*P < .05; **P < .01; ***P < .001). The detection limit for the assay is represented in black dotted line (10 CFU).

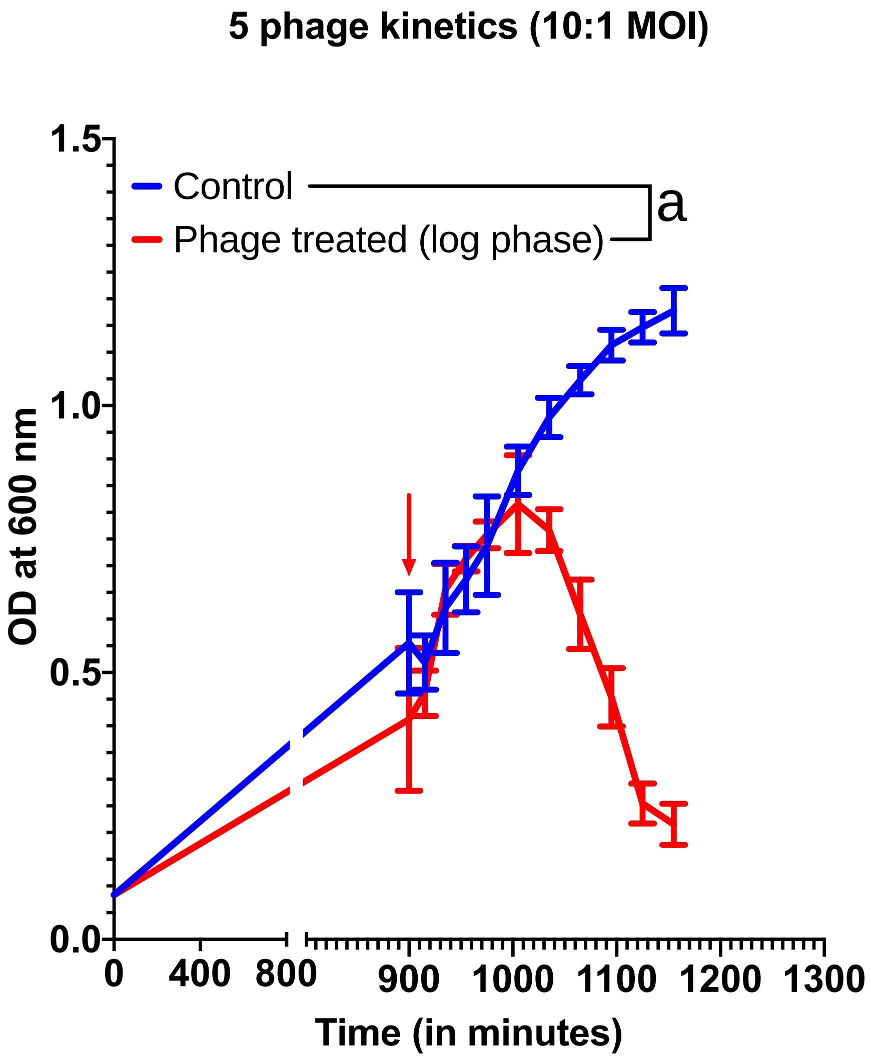


**Supplementary Figure S4. Phage cocktail (D29, TM4, Che7, PDRPv and PDRPxv) kinetics at 10 MOI**. Red arrow represents the time point phage solution was added to the *M. smegmatis* cultures. Multiple t-test with Holm-Sidak method was used to estimate the statistical significance ($p<0.05$). $\alpha$ represents a significant difference ($p<0.05$) between control and five-phage cocktail (D29, TM4, Che7, PDRPv and PDRPxv) treated (log phase) for all time points between 1080 min and 1200 min.

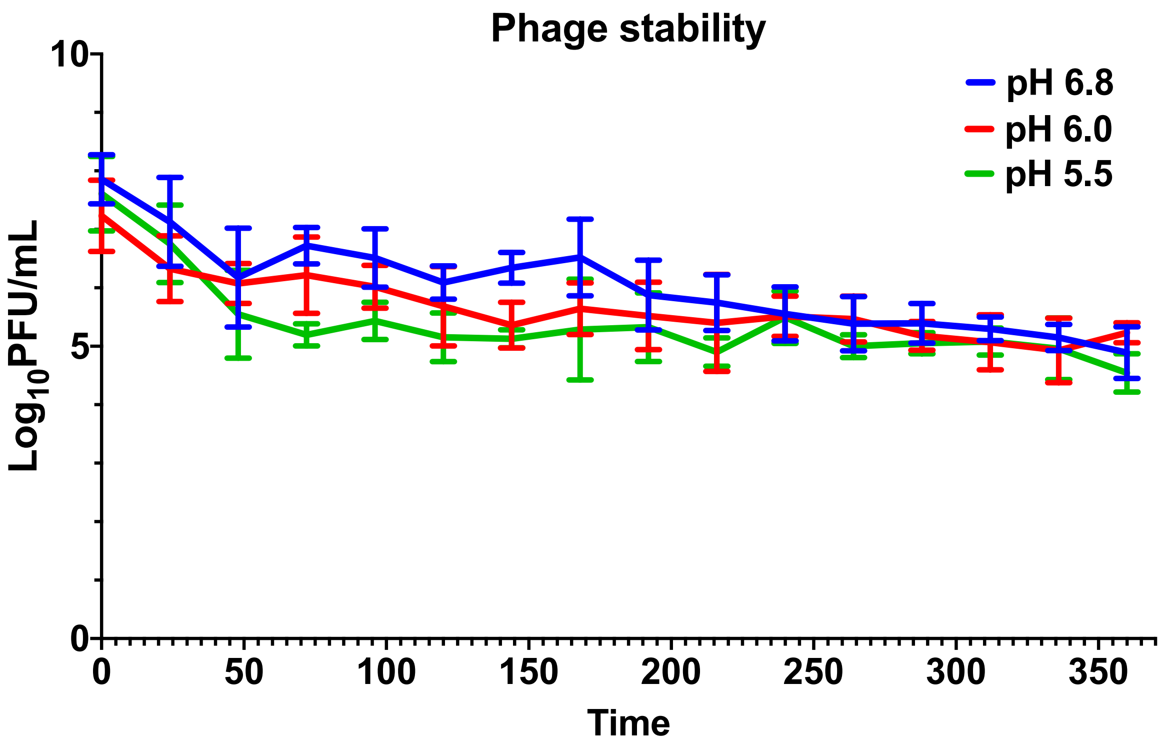


**Supplementary Figure S5. Stability of phages at various pH in 7H9 medium over time.**

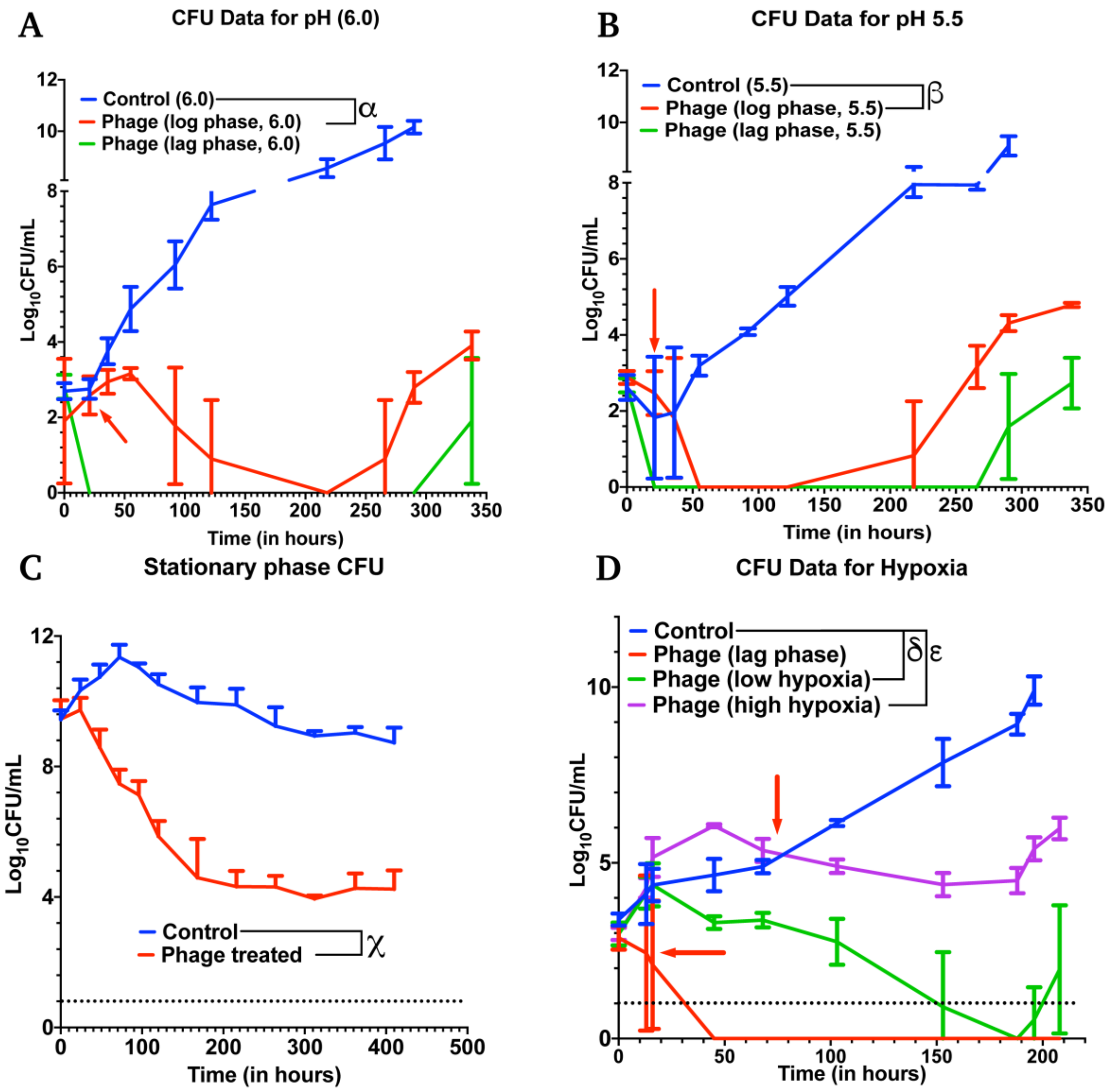


**Supplementary figure S6. Mycobacteriophages are effective under various disease-mimicking conditions.** Colony Forming Unit (CFU) measurements over time of *M. smegmatis* cultures treated with a 5-phage cocktail at **A)** pH condition of 6.0, **B)** pH condition of 5.5, **C)** stationary phase, and **D)** hypoxic conditions. Red arrow represents the time point phage solution was added to the cultures treated in the log phase. For **A**, **B**, and **C**, multiple t-test with Holm-Sidak method was used to estimate the statistical significance. α represents a significant difference ($p<0.05$) between control and phage treated (pH 6.0, log phase) at all time points between 55 h and 290 h. β represents a significant difference ($p<0.05$) between control and phage treated (pH 5.5, log phase) at all time points between 55 h and 290 h. χ represents a significant difference ($p<0.05$) between control and phage treated (stationary phase) at all time points between 48 h and 410 h. For **D,** 2-way ANOVA with Sidak's multiple comparisons test was used ($p<0.05$). δ represents a significant difference between control and phage (at low hypoxia) at all time points between 45 h and 196 h. ε represents a significant difference between control and phage (at high hypoxia) at all time points between 103 h and 196 h. The detection limits for the assays are represented in black dotted line (10 CFU).

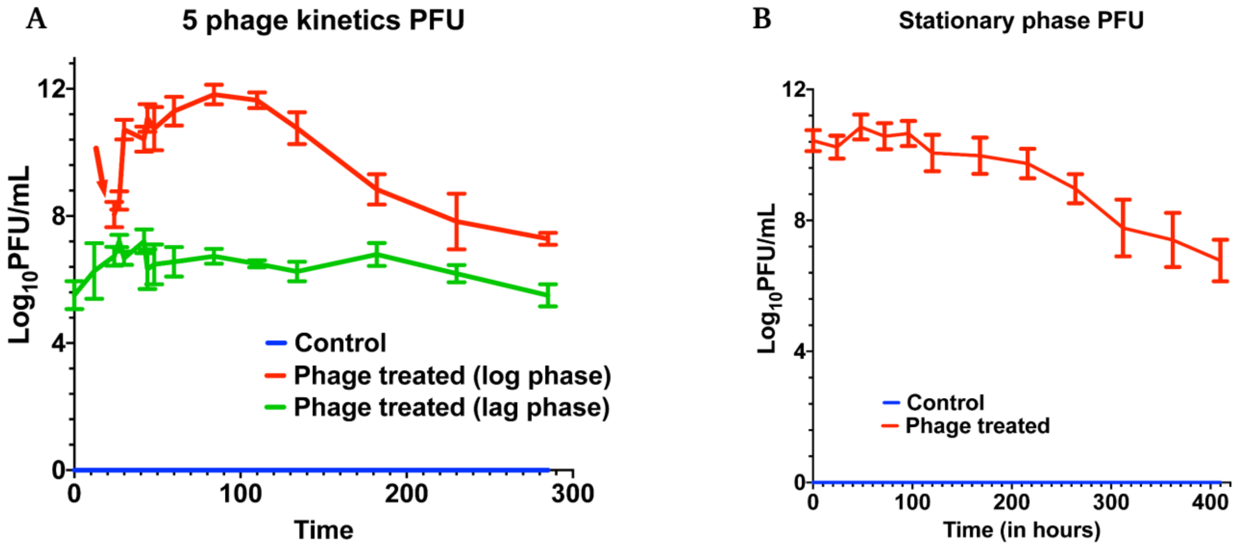


**Supplementary figure S7. Phages amplify and persist throughout the duration of the experiment.** Plaque Forming Unit (CFU) measurements over time of M. smegmatis **A)** lag and log phase cultures treated with 5-phage cocktail and **B)** stationary phase cultures treated with 5-phage cocktail. Red arrow represents the time point phage solution was added to the cultures treated in the log phase.

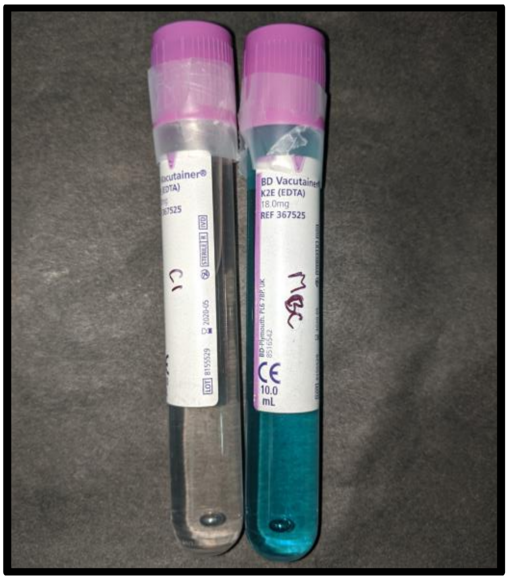


**Supplementary figure S8. Methylene Blue as an indicator for hypoxia**. Photograph of vaccutainers with *M. smegmatis* culture. On the left, an overnight grown *M. smegmatis* culture. Methylene Blue discolouration indicates that the culture has reached hypoxia. On the right, media after overnight incubation without bacteria where no discolouration is observed.

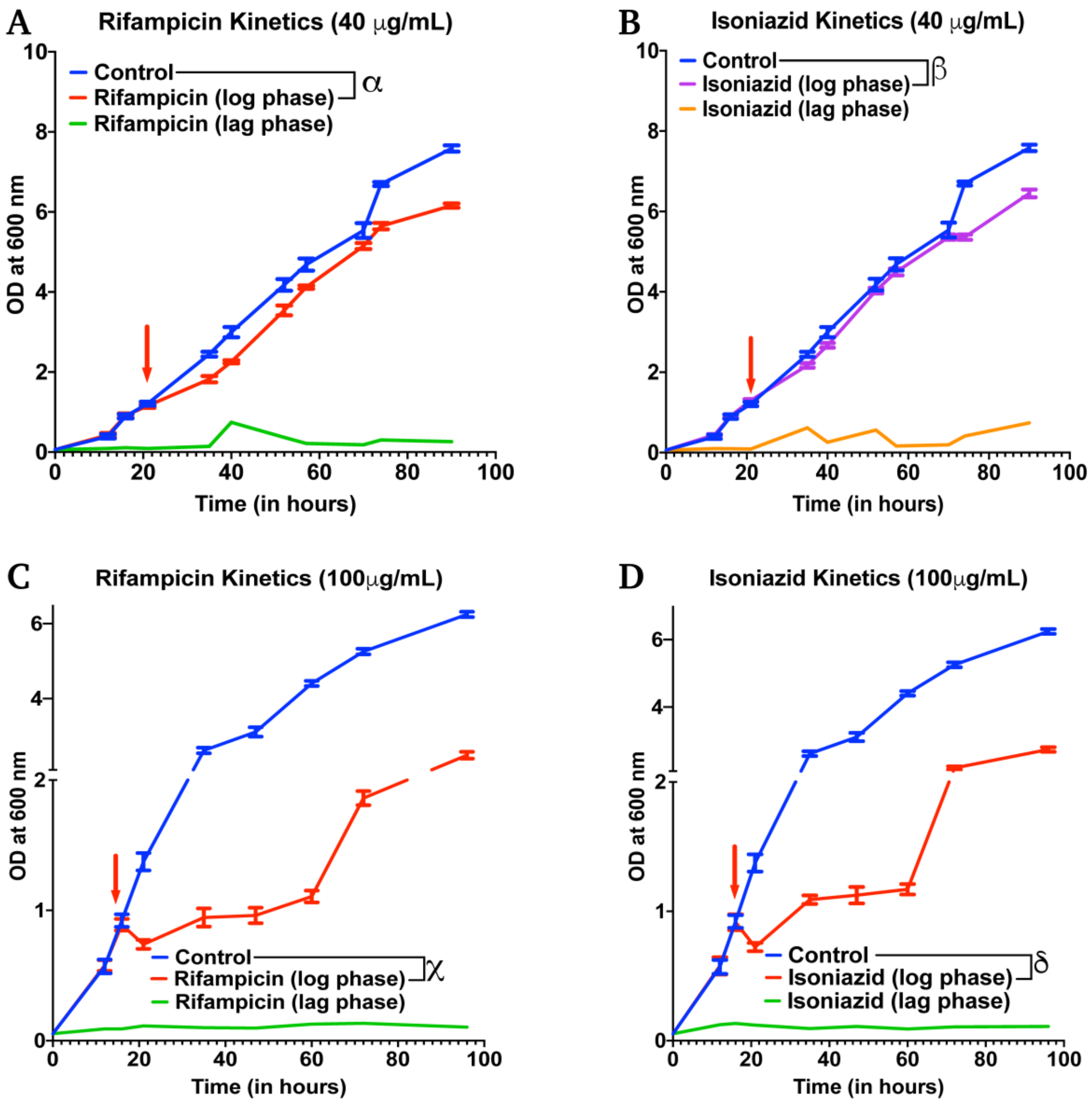


**Supplementary figure S9. Growth kinetics of *M. smegmatis* cultures under antibiotic treatments.** Growth kinetics of *M. smegmatis cultures* treated with **A)** rifampicin (40 µg/mL) - α represents a significant difference ($p<0.05$) between control and rifampicin treated (log phase) at all time points between 35 h and 90 h, excluding 70 h, **B)** isoniazid (40 µg/mL)- β represents a significant difference ($p<0.05$) between control and isoniazid treated (log phase) at all, **C)** rifampicin (100 µg/mL)- χ represents a significant difference ($p<0.05$) between control and rifampicin treated (log phase) for all time points between 21 h and 96 h, **D)** isoniazid (100 µg/mL)- δ represents a significant difference ($p<0.05$) between control and isoniazid treated (log phase) for all time points between 21 h and 96 h. Red arrow represents the time point antibiotic was added to the cultures. time points between 74 h and 90 h.

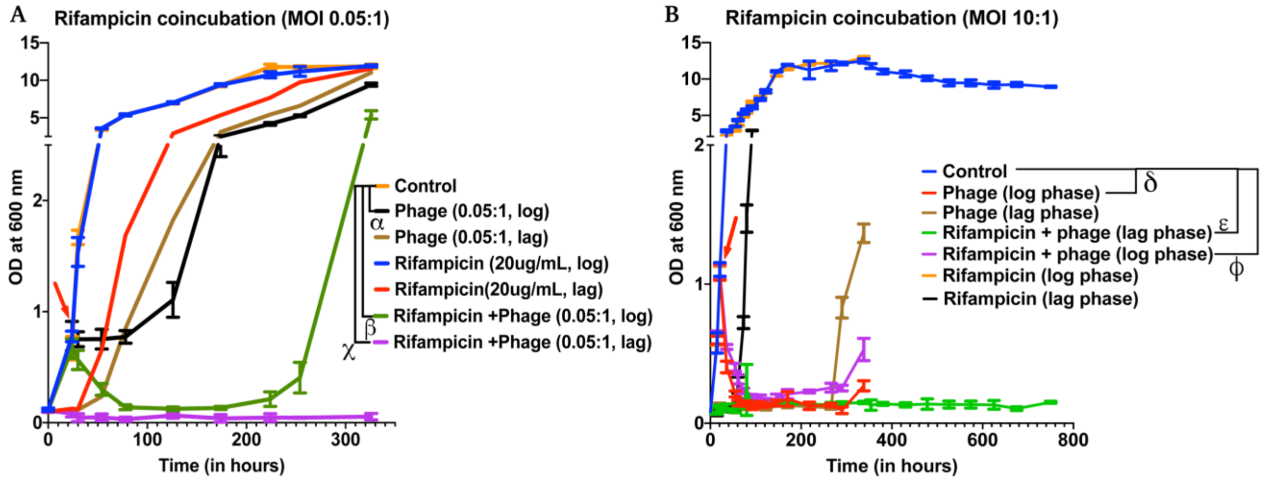


**Supplementary figure S10. Rifampicin (20 µg/mL) and phage coincubation**. **A)** at 5-phage cocktail MOI of 0.05- α**, and** β represent a significant difference between control and phage treated (log phase) or rifampicin + phage (log phase) for all time points between 30 h and 326 h. χ represents a significant difference between control and rifampicin + phage (lag phase) for all time points between 24 h and 326 h. **B)** at 5-phage cocktail MOI of 10- δ represents a significant difference ($p<0.05$) between control and phage cocktail treated (log phase) at all time points between 36 h and 338 h. ε represents a significant difference ($p<0.05$) between control and rifampicin + phage treated (lag phase) at all time points between 15 h and 750 h. ϕ represents a significant difference ($p<0.05$) between control and rifampicin + phage treated (log phase) for all time points between 36 h and 338 h. Red arrow represents the time point phage+ rifampicin solution was added to the cultures treated at log phase. For A, 2-way ANOVA with Sidak's multiple comparisons test was used to determine statistical significance ($p<0.05$). For B, multiple t-test with Holm-Sidak method was used to estimate the statistical significance ($p<0.05$).

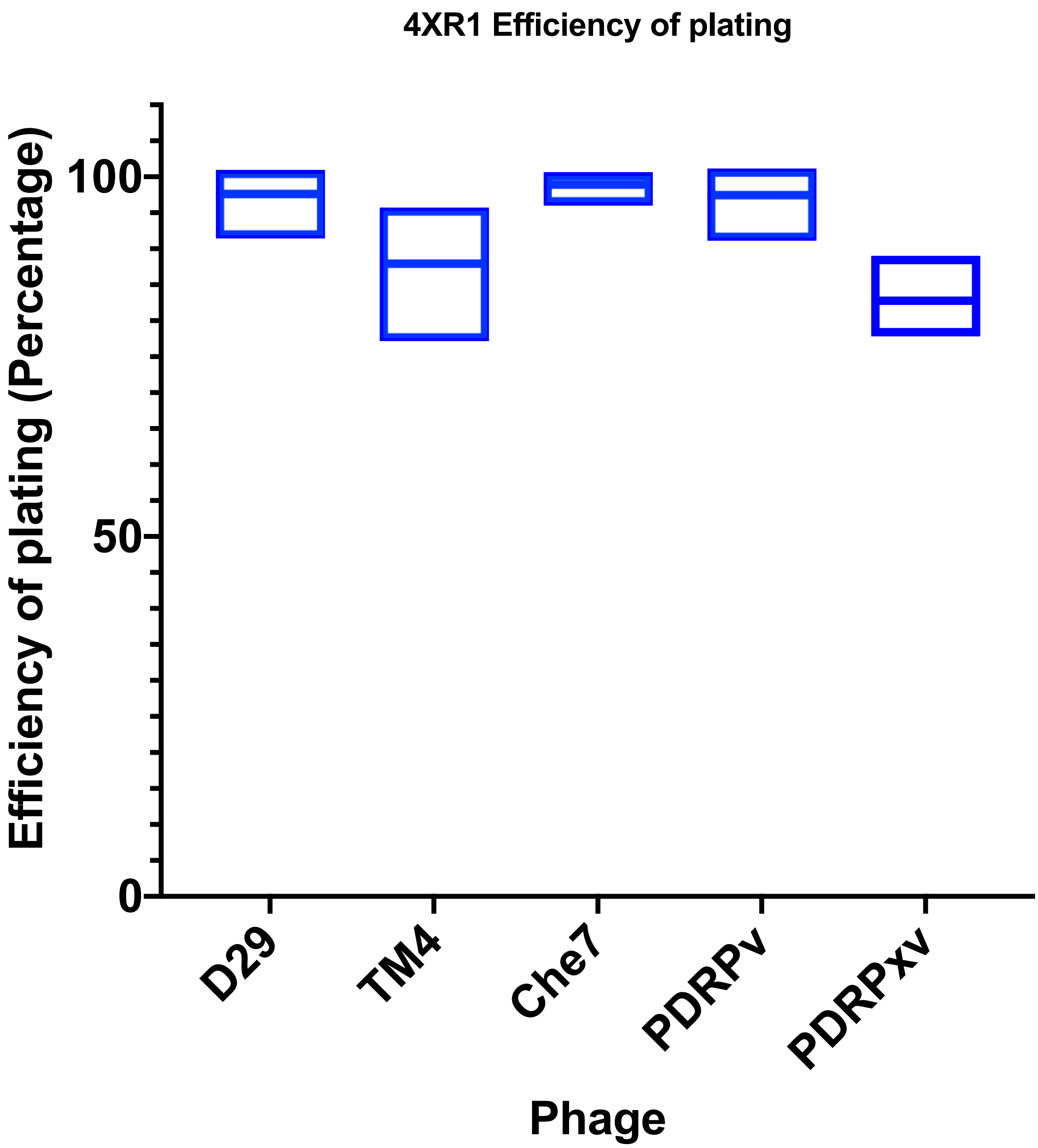


**Supplementary Figure S11. Phage efficiency of plating for isoniazid resistant M. smegmatis (4XR1 strain)**. Experiments were carried out in triplicates (n=3). Values are normalized against phage titers against *M. smegmatis* mc$^2$ 155.
